# Stochastic comparison of synchronization in activator- and repressor-based coupled gene oscillators

**DOI:** 10.1101/2021.07.08.451708

**Authors:** A B M Shamim Ul Hasan, Supravat Dey, Hiroyuki Kurata, Abhyudai Singh

## Abstract

Inside living cells, proteins or mRNA can show oscillations even without a periodic driving force. Such genetic oscillations are precise timekeepers for cell-cycle regulations, pattern formation during embryonic development in higher animals, and daily cycle maintenance in most organisms. The synchronization between oscillations in adjacent cells happens via intercellular coupling, which is essential for periodic segmentation formation in vertebrates and other biological processes. While molecular mechanisms of generating sustained oscillations are quite well understood, how do simple intercellular coupling produces robust synchronizations are still poorly understood? To address this question, we investigate two models of coupled gene oscillators - activator-based coupled oscillators (ACO) and repressor-based coupled oscillators (RCO) models. In our study, a single autonomous oscillator (that operates in a single cell) is based on a negative feedback, which is delayed by intracellular dynamics of an intermediate species. For the ACO model (RCO), the repressor protein of one cell activates (represses) the production of another protein in the neighbouring cell after a intercellular time delay. We investigate the coupled models in the presence of intrinsic noise due to the inherent stochasticity of the biochemical reactions. We analyze the collective oscillations from stochastic trajectories in the presence and absence of explicit coupling delay and make careful comparison between two models. Our results show no clear synchronizations in the ACO model when the coupling time delay is zero. However, a non-zero coupling delay can lead to anti-phase synchronizations in ACO. Interestingly, the RCO model shows robust in-phase synchronizations in the presence and absence of the coupling time delay. Our results suggest that the naturally occurring intercellular couplings might be based on repression rather than activation where in-phase synchronization is crucial.

## 1. INTRODUCTION

The phenomena of oscillations are widely found in living systems. At the cellular level, the gene products such as mRNA/proteins can show sustained oscillations even without any periodic driving force Goldbeter and Berridge (1996); Novák and Tyson (2008); Forger (2017). Such gene oscillations are found in various contexts and across organisms. The popular examples of genetic oscillators are the circadian clock Stokkan et al. (2001); Ray et al. (2020), cell-cycle clock Hara et al. (1980); Pomerening et al. (2003), and segmentation clock Palmeirim et al. (1997); Zinani et al. (2021). The circadian clocks are responsible for maintaining diurnal cycle in most organisms, cell-cycle clocks regulate precise time in cell division, and segmentation clocks dictate accurate rhythmic somite formation during embryonic development in vertebrates.

For the generation of sustained gene oscillations, there are nonlinear regulatory circuits Novák and Tyson (2008). Depending on contexts, the complexity of such a circuit may vary. However, the basic regulatory motif for many gene oscillators, including the circadian clock, cell-cycle clock, and segmentation clock, is time-delayed negative feedback. The time-delays can arise due to the dynamics of intermediates before the final repression of the clock genes. In eukaryotes, the most contribution of the timedelay comes from the mRNA exportation from the cell nucleus to cytoplasm, importation of cytoplasmic protein to the cell nucleus where transcriptional repression happens, and post-transcriptional modification dynamics. Theoretical models incorporate such delays using explicitly Lewis (2003) or incorporating the dynamics of several intermediates Griffith (1968); Morelli and Jülicher (2007). Such gene regulatory circuits are subject to unavoidable fluctuations due to the inherent stochasticity of biochemical reactions occurring in low molecular abundance Elowitz et al. (2002); Shaffer et al. (2017). Amid such noise, cells produce gene oscillations that are usually remarkably precise Geva-Zatorsky et al. (2006); Webb et al. (2016); Keskin et al. (2018). Researchers also successfully engineered synthetic circuits based on delayed repression to generate robust and persistent oscillations in bacteria Elowitz and Leibler (2000); Potvin-Trottier et al. (2016) and mammal cells Tigges et al. (2009).

Although individual cells can act as autonomous oscillators, they must oscillate collectively for proper functions in many biological processes. For example, in the case of segmentation clocks, adjacent cells oscillate in synchrony to produce accurate vertebra formation Lewis (2003); Venzin and Oates (2020). To create such collective oscillations, intercellular coupling between neighbouring cells is essential. Such coupling often involves complex signaling pathways, and its role in synchronization is not well understood ?. Effectively, via intercellular coupling, dynamical changes in molecular concentration of relevant proteins in a cell affect the production of the clock genes in the neighbouring cells. The process of a cell responding to the signal from the neighbours could involve significant time. In essence, the intercellular coupling can activate or repress the production from clock genes in neighbouring cells with some time delay. In this paper, we aim to understand its role by studying two simple models – activator-based coupled oscillators (ACO), and repressor-based coupled oscillators (RCO) models in the presence of coupling delay. For our study we consider, the single autonomous oscillator is based on delayed inhibition where the intracellular time delay is caused by the dynamics of an intermediate species.

## 2. MODEL FORMULATION

### 2.1 Dynamics of single oscillator

We first discuss the dynamics of the single autonomous oscillator. For this study, we consider a general model for gene oscillator based on delayed inhibition, consisting of three proteins X, Y, and Z (see Fig.1A). The protein X activates Y, and Y activates the expression of Z protein.

**Fig. 1.**
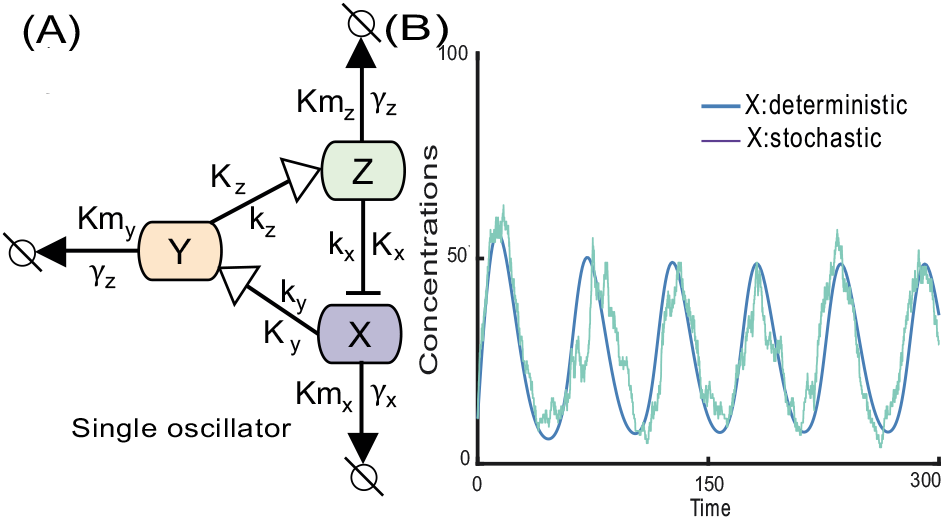
The schematic of single gene oscillator. (A) A negative feedback loop with at three components Nováak and Tyson (2008); where X → Y means ‘X activates Y’ and X⊣Y means ‘X inhibits Y’ (B) Trajectories of the deterministic and stochastic simulation of the production of proteins X with *n* = 2. Other parameters: *k_x_* = *k_y_* = *k_z_* = 10, *γ_x_*=*γ_y_*=*γ_z_*=8, *K_x_*=*K_y_*=*K_z_*=30, *Km_x_*=*Km_y_*=*Km_z_*=30.

Finally, protein Z represses the X productions and closes the negative feedback. The dynamics of Y and Z cause an effective time delay for the repression. The deterministic dynamics of the concentration of species (denoted as corresponding lower case letters *x, y*, and *z*) are given by:

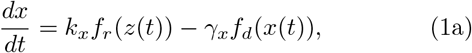

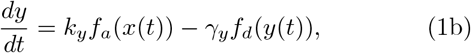

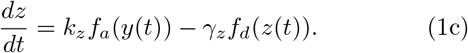

The parameter *k_x_, k_y_*, and *k_z_* are the maximum synthesis rates for the protein X, Y and Z, respectively. The maximum degradation rates are denoted by symbols *γ_x_, γ_y_*, and *γ_z_*. The functions *f_r_* and *f_a_* are the nonlinear Hill-type functions for the repression and activation Alon (2011); Wilhelmováa (1996). These functions are given by

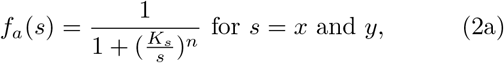

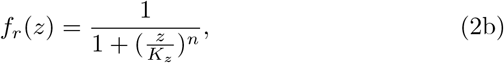

where *n* is the Hill coefficient for the activation and repression, and *K_x_, K_y_* and *K_z_* the dissociation constant, representing the values of respective protein concentrations where repression and activation become half of its maximum value. We consider the Michaelis-Menten type degradation, assuming that substrate concentrations are in excess and association equilibria quickly attained. The degradation function *f_d_* for a species S is given by

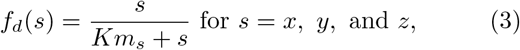

where the parameter *Km_s_* dissociation constant corresponding to Michaelis-Menten degradation.

In circadian clocks, protein degradation is controlled by phosphorylation, ubiquitination, and proteasomal degradation and thus it is reasonable to assume Michaelis–Menten kinetics Goldbeter (2013). Therefore, we used the Michaelis-Menten degradation of protein Gonze et al. (2005); Griffith (1968); Purcell et al. (2010); Novák and Tyson (2008). We note that in the limit of linear activation and linear degradation, the model reduced to the well-known Goodwin oscillator, where the repression term is the only source of nonlinearity Goodwin (1965); Griffith (1968). For the latter case, to get sustained oscillations, the Hill coefficient for the repression must be greater than 8 Griffith (1968). As the Goodwin oscillator assumes a minimal regulatory mechanism for generating sustained oscillations, it is beneficial for mathematical insights Griffith (1968); Dey and Singh (2020). However, such a high Hill coefficient value is often considered biologically unrealistic Gonze and Abou-Jaoudáe (2013). With other nonlinearities coming from activation and Michaelis-Menten, in our model, one can get sustained oscillations even with the Hill coefficient *n* = 2. Here, the typical trajectories of species X for *n* = 2 is shown in Fig. 1B.

### 2.2 Dynamics of coupled oscillators

The oscillator in each cells are coupled to its adjacent cell via signaling pathway. Such coupling are essential for synchronization between oscillations in neighboring cells.

The signaling pathways often complex that involves several molecular species undergoing many biochemical reactions. Such complexity makes it challenging for theoretical studies of synchronizations Venzin and Oates (2020).

Here we study two different models of two coupled oscillators with opposite coupling mechanisms, schematically shown in Fig. 2. In our models, we do not have additional signaling species for intercellular coupling. The protein Z itself acts as a signaling molecule and affects the production of the protein X in the other cell with some time delay *τ*. We consider two coupling mechanisms: activator-based coupled oscillators (ACO) and repressor-based coupled oscillators (RCO). More specifically, in ACO, the protein Z in a cell activates the production of the protein X in the other cell. Whereas in RCO, the protein Z in a cell represses the production of the protein X in the other. In our study, we assume both oscillators are identical and the coupling acts symmetrically between two oscillators. The coupling directly affects the production of protein X. The deterministic dynamics of protein X concentration for a cell *i* = {1, 2} is

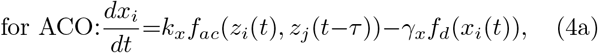

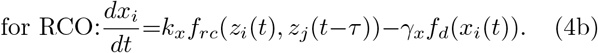

where *j* is the neighbour of *i*. The coupling functions for ACO *f_ac_* and for RCO *f_rc_* are

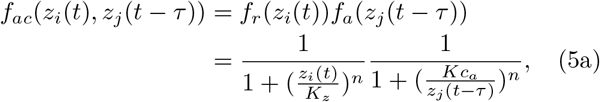

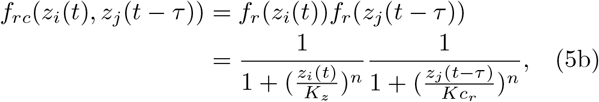

where *Kc_a_* and *Kc_r_* are the corresponding coupling dissociation constants. We note that for ACO, in the limit of *Kc_a_* → 0, both oscillators become uncoupled, whereas for RCO the uncoupled limit is *Kc_r_* → ∞. The dynamics of Y and Z proteins are not directly affected by the coupling and are given by

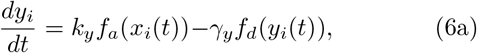

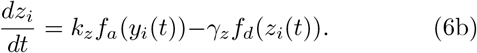

**Fig. 2.**
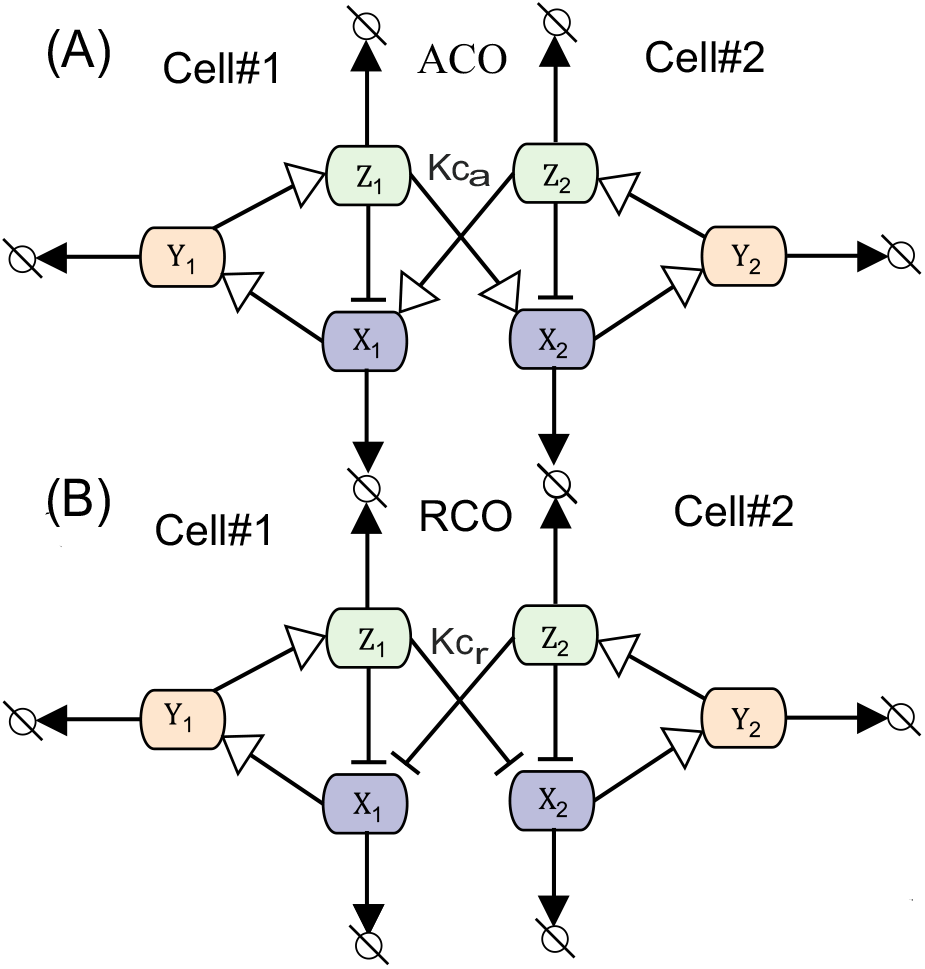
The Schematic diagrams of the models of coupling oscillators. A single oscillator operates in a single cell. Both cells are identical and the coupling acts symmetrically between them. (A) The ACO model: the protein Z in one cell activates the production of protein X in the other cell. (B) The ACO model: the protein Z in one cell represses the production of protein X in the other cell.

The gene expression is noisy as the biochemical reactions are inherently stochastic. As the molecular species inside cells present in low abundances where such noise become play important role in the dynamics. We study the above coupled systems in the presence of such intrinsic noise. As our coupling explicitly depends on the time-delay, we can not use traditional stochastic simulations algorithm Gillespie (1977). Here, we employ the time-delay stochastic simulation algorithm as proposed in Barrio et al. (2006).

## 3. RESULTS

### 3.1 Collective oscillations without intercellular time delay

First, we study collective oscillations in the absence of intercellular coupling time delay (*τ* = 0) by varying the coupling dissociation constant which is a measure of coupling strength. Then, we study the effect of the coupling time delay. As the ACO and RCO models work very differently, comparing their the collective oscillations in these two models can be tricky. We do the following mathematical comparison between the ACO and RCO.

#### Mathematical comparison between ACO and RCO

It is important to identify different structures between the competitive models and to understand some characteristic structures within each model. This is very much in the spirit of the mathematically controlled comparison Hasan and Kurata (2017); Hasan et al. (2019); Hasan and Kurata (2016); Alves and Savageau (2000). To set a sound basis for comparison between two competitive models, the equivalence between them should be guaranteed with respect to their function and corresponding kinetics. Therefore, we conserve the other functions between them, to compare a specific function of two competitive models, while reducing the search space by setting the corresponding kinetic parameters to the same values.

#### A. Coupled oscillators with conserved expression levels

To conserve the expression levels of protein X in the ACO and RCO models, we use the parameter set presented in Table. 2. We only vary the value of the coupling dissociation constants, while the values of the other kinetic parameters within each model and between the competitive models are the same. The results are presented in Fig. 3. The RCO model shows robust in-phase synchronization. Whereas, the ACO model does not show any clear phase synchronizations – cell-1 and cell-2 seems to oscillate randomly.

**Table 1.** Definition of the kinetic parameters

| Kinetic parameter | Description |
| --- | --- |
| $k_x, k_y, k_z$ | synthesis rate constant |
| $\gamma_x, \gamma_y, \gamma_z$ | protease (enzyme) rate constant |
| $K_x, K_y, K_z$ | disassociation const. for activation/repression |
| $Km_x, Km_y, Km_z$ | disassociation const. for Michaelis-Menten |
| $Kc_a, Kc_r$ | coupling disassociation constant |
| $n$ | Hill coefficient |
| $\tau$ | coupling time-delay |

**Table 2.** Parameters for the ACO and RCO models to conserve the expression levels.

| Each network | ACO | RCO |
| --- | --- | --- |
| Corresponding parameters | $k_x=k_y=k_z=10$<br>$\gamma_x=\gamma_y=\gamma_z=8$<br>$K_x=K_y=30$<br>$K_z=30$<br>$Km_x=Km_y=30$<br>$Km_z=30$<br>$Kc_a=1.1$<br>$n=3$<br>$\tau=0$ | $k_x=k_y=k_z=10$<br>$\gamma_x=\gamma_y=\gamma_z=8$<br>$K_x=K_y=30$<br>$K_z=30$<br>$Km_x=Km_y=30$<br>$Km_z=30$<br>$Kc_r=30$<br>$n=3$<br>$\tau=0$ |

**Fig. 3.**
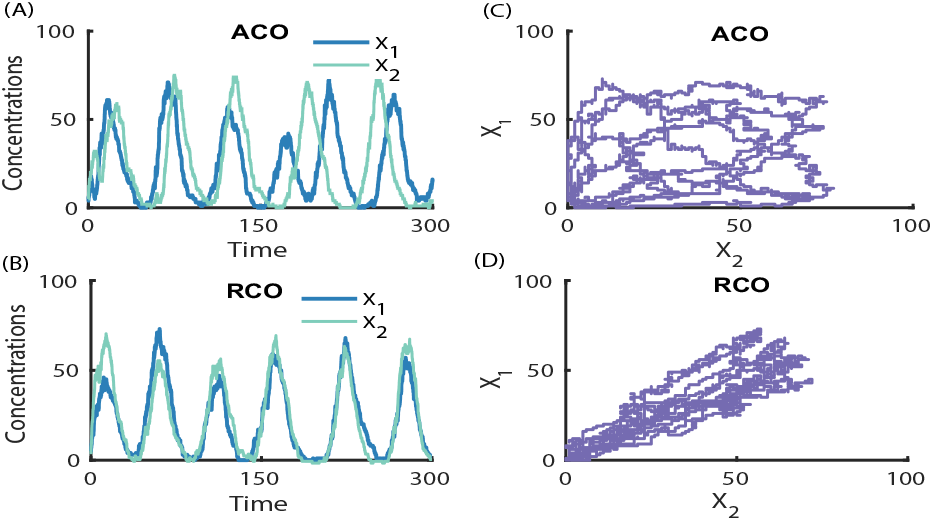
Stochastic simulation results for the ACO and RCO models at conserved expression levels. Trajectories of concentration of protein X for cells 1 and 2 for the ACO and RCO models are plotted in (A) and (B), respectively. (C) and (D) are the scatter plots of X_1_ and X_2_ corresponding to (A) and (B). For the RCO, the trajectories of both cells follow each other, suggesting a strong in-phase synchronization. For the ACO, the trajectories of both cells seem random, suggesting no clear phase synchronization.

#### B. Coupled oscillators with conserved period

Is the collective oscillation observed in the conserved expression comparison also holds across other parameters? Here, we make another comparison by keeping the period of the oscillations in both the model the same. The parameters for this case are presented in Table. 3. We only vary the value of the coupling dissociation constants, while the values of the other kinetic parameters within each model and between the competitive models are the same. The results are presented in Fig. 4. As in the conserved expression comparison case, we find robust in-phase synchronization for the RCO, while the ACO model shows no clear phase synchronizations.

**Table 3.** Parameters for the ACO and RCO models to conserve the expression period.

| Each network | ACO | RCO |
| --- | --- | --- |
| Corresponding parameters | $k_x=k_y=k_z=10$<br>$\gamma_x=\gamma_y=\gamma_z=8$<br>$K_x=K_y=13.55$<br>$K_z=13.55$<br>$Km_x=Km_y=13.55$<br>$Km_z=13.55$<br>$Kc_a=1.1$<br>$n=3$<br>$\tau=0$ | $k_x=k_y=k_z=10$<br>$\gamma_x=\gamma_y=\gamma_z=8$<br>$K_x=K_y=13.55$<br>$K_z=13.55$<br>$Km_x=Km_y=13.55$<br>$Km_z=13.55$<br>$Kc_r=30$<br>$n=3$<br>$\tau=0$ |

**Fig. 4.**
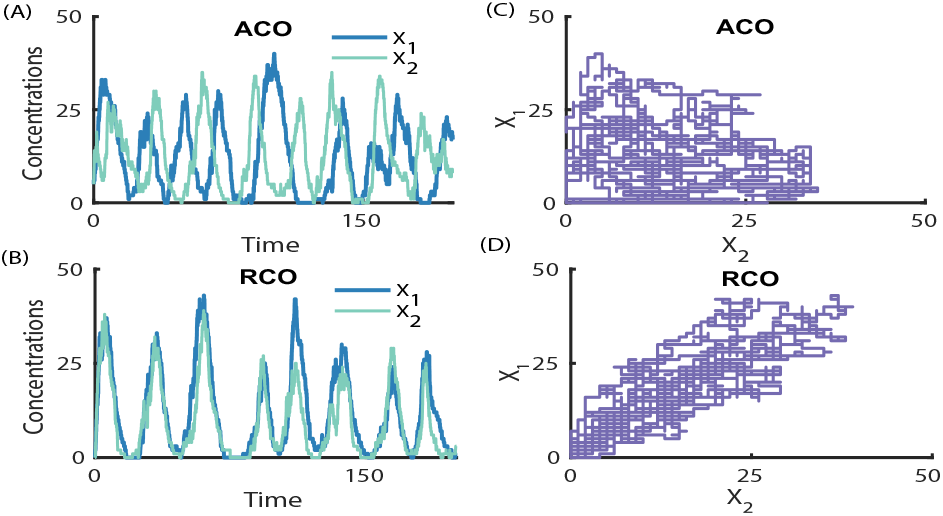
Stochastic simulation results for the ACO and RCO models at conserved period. Trajectories of concentration of protein X for cells 1 and 2 for the ACO and RCO models are plotted in (A) and (B), respectively. (C) and (D) are the scatter plots of X_1_ and X_2_ corresponding to (A) and (B). For the RCO, the trajectories of protein X for both cells show strong in-phase synchronization. For the ACO, the trajectories of both cells seem random.

We also run several stochastic simulations by perturbing parameters presented in Table 2 and Table 3. We find the robust in-phase synchronization for the RCO across parameters. Again, no clear phase synchronization is observed in the ACO model.

### 3.2 Effect of intercellular time delay

Now, we turn our focus on the effect of the intercellular coupling time delay. Time delay can have an interesting effects on the coupled oscillations Kim et al. (2010); Joshi et al. (2020); Rashid and Kurata (2020); Amdaoud et al. (2007). It also plays a critical role in synchronization between the coupled feedback oscillators Smolen et al. (2001). For example, it can lead to alternative in-phase synchronization and anti-phase as the delay time is varied Jörg et al. (2018); Giordano et al. (2019); Dey et al. (2020). Here, we quantify the collective oscillation by the Pearson correlation coefficient in the presence and absence of coupling delay time *τ*. If *X*_1_(*t*) and *X*_2_(*t*) are the stochastic variables representing the concentration protein X in cell 1 and 2 at time *t*, respectively, then the Pearson correlation, *R* between protein X level in two cells is given by,

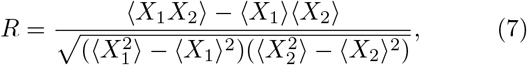

where the angular bracket 〈·〉 denotes the average over time and ensembles. The value of correlation *R* is 1 if protein X in both cells oscillate in perfect in-phase, –1 if both cells oscillate in perfect anti-phase, 0 if both cells oscillate randomly. We measure the correlation coefficient *R*, for coupling delay *τ* = 0 and *τ* = 12. The period of the oscillations is 24 when *τ* = 0. For a given *τ*, we compute R for different values of coupling dissociation constant *Kc_a_* or *Kc_r_*. Each R value is computed averaging over hundreds of long trajectories with trajectory length 8000.

In Fig. 5(A), we plot R as a function of *Kc_a_* and *Kc_r_* for the ACO and RCO model. For *τ* = 0, *R* remains close to zero throughout for the ACO, whereas R becomes close to 1 for large values of *Kc_r_*, showing in-phase synchronizations in the RCO and no synchronization the in ACO model. This consistent with the observation discussed in the above section. The RCO continues to show in-phase synchronization in the presence of time delay. Interestingly, for a non-zero *τ*, the value of R becomes close to −1 for large dissociation constant for the ACO, suggesting the development of anti-phase synchronization in the presence of time delay. The in-phase synchronization in the RCO and anti-phase synchronization in the ACO for a finite *τ* can also be clearly seen from the stochastic trajectory plots (Fig. 5(B) and (C)).

**Fig. 5.**
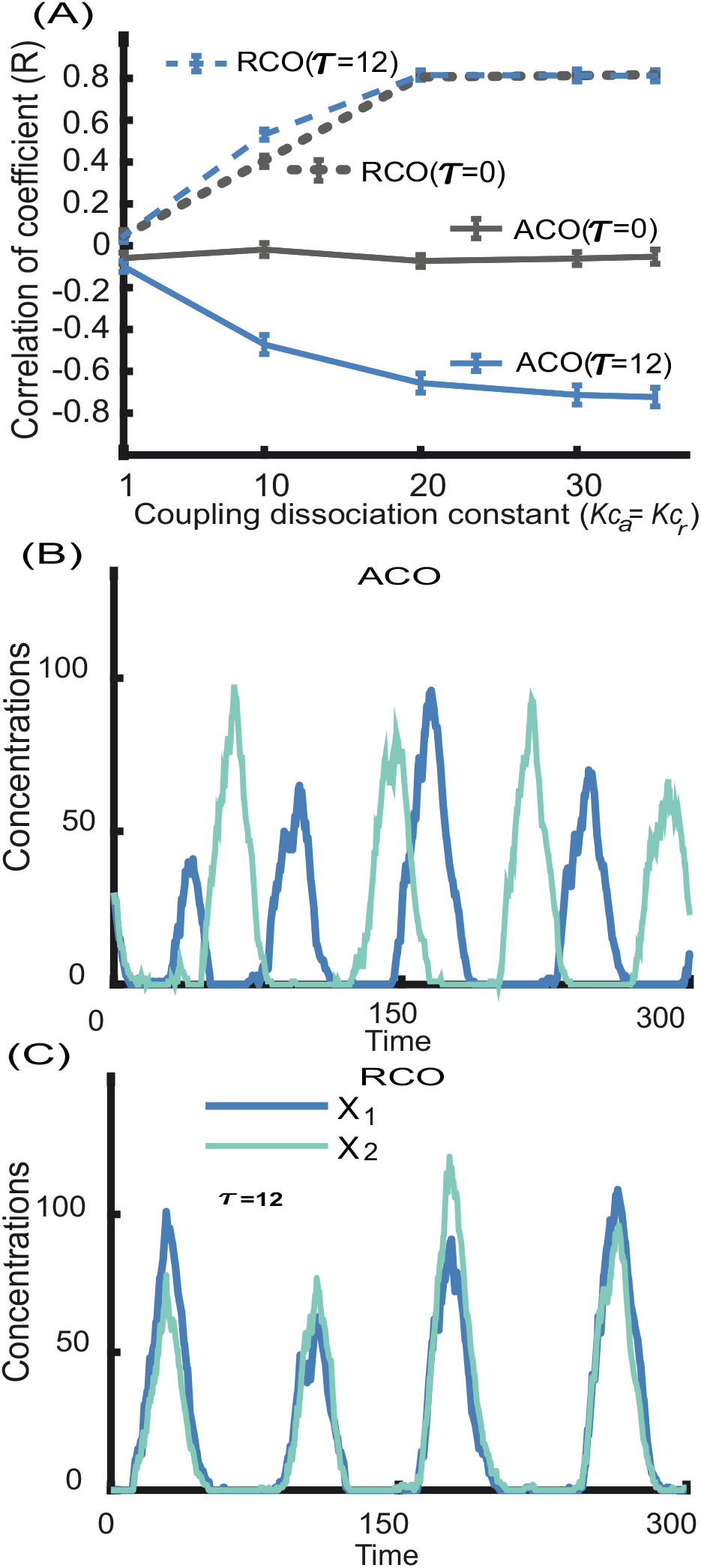
Effect of coupling time-delay. (A) The correlation *R* is plotted for *τ* = 0 and *τ* = 12 as a functions the coupling dissociation constant. For the RCO, R approaches to 0.8 irrespective of the value of *τ*, showing strong in-phase synchronizations. While for the ACO, the synchronization behavior depends on the value of *τ*. When *τ* = 0, two cells oscillates randomly results R ≈ 0. For *τ* = 0, *R* ≈ −0.8 for large *Kc_*a*_*, suggesting an anti-phase synchronization. (B) and (C) The stochastic trajectories for the ACO and RCO for *τ* =12 for *Kc_r_* = 30, *Kc_a_* = 1.1. Other parameters are shown Table 3.

## 4. CONCLUSION

Synchronizations of genetic oscillators are crucial for many biological processes, including precise vertebra formation during embryonic development Lewis (2003); Venzin and Oates (2020). Amid inherent stochastic fluctuations due to biochemical reactions, how do the intercellular coupling leading to robust synchronizations is poorly understood? To understand the role of intercellular coupling, we have developed two models for coupling, namely the ACO (activator-based coupled gene oscillator) and RCO (repressor-based coupled gene oscillators). The impacts of signaling molecules are effectively incorporated via the usual Hill functions Alon (2011); Wilhelmová (1996). We have included an effective coupling delay due to the dynamics of signaling molecules. The single autonomous oscillator in our study is based on delayed inhibition where the dynamics of intermediate species cause the delay.

We have solved our stochastic systems of coupled oscillators in the presence and absence of delay using the algorithm as proposed in Barrio et al. (2006). We have found that the RCO show robust in the presence and absence of time delay. Interestingly, in the ACO, we have not found a clear synchronization without coupling delay. However, in the presence of coupling delay, the ACO can lead to anti-phase synchronization. Our results suggest that the naturally occurring intercellular couplings might be based on repression rather than activation, where inphase synchronization is crucial.

## ACKNOWLEDGEMENTS

AS is supported by the National Science Foundation grant IR01GM124446-01 and the ARO grant W911NF-19-1-0243.

